# LogoMotif: a comprehensive database of transcription factor binding site profiles in Actinobacteria

**DOI:** 10.1101/2024.02.28.582527

**Authors:** Hannah E. Augustijn, Dimitris Karapliafis, Kristy Joosten, Sébastien Rigali, Gilles P. van Wezel, Marnix H. Medema

## Abstract

Actinobacteria undergo a complex multicellular life cycle and produce a wide range of specialized metabolites, including the majority of the antibiotics. These biological processes are controlled by intricate regulatory pathways, and to better understand how they are controlled we need to augment our insights into the transcription factor binding sites. Here, we present LogoMotif (https://logomotif.bioinformatics.nl), an open-source database for characterized and predicted transcription factor binding sites in Actinobacteria, along with their cognate position weight matrices and hidden Markov models. Genome-wide predictions of binding site locations in *Streptomyces* model organisms are supplied and visualized in interactive regulatory networks. In the web interface, users can freely access, download and investigate the underlying data. With this curated collection of actinobacterial regulatory interactions, LogoMotif serves as a basis for binding site predictions, thus providing users with clues on how to elicit the expression of genes of interest and guide genome mining efforts.

**Highlights:**

- Actinobacterial regulatory networks are key for compound discovery, including antibiotics.
- Contains ∼400 validated and ∼12,100 predicted interactions, presented in interactive networks.
- Serves as foundation for regulatory predictions in the gene cluster detection tool, antiSMASH.
- LogoMotif’s data and algorithms provide knowledge on expression and functional inference of genes.
- LogoMotif aids in the discovery of novel chemistry within Actinobacteria and beyond.

**Graphical abstract:** 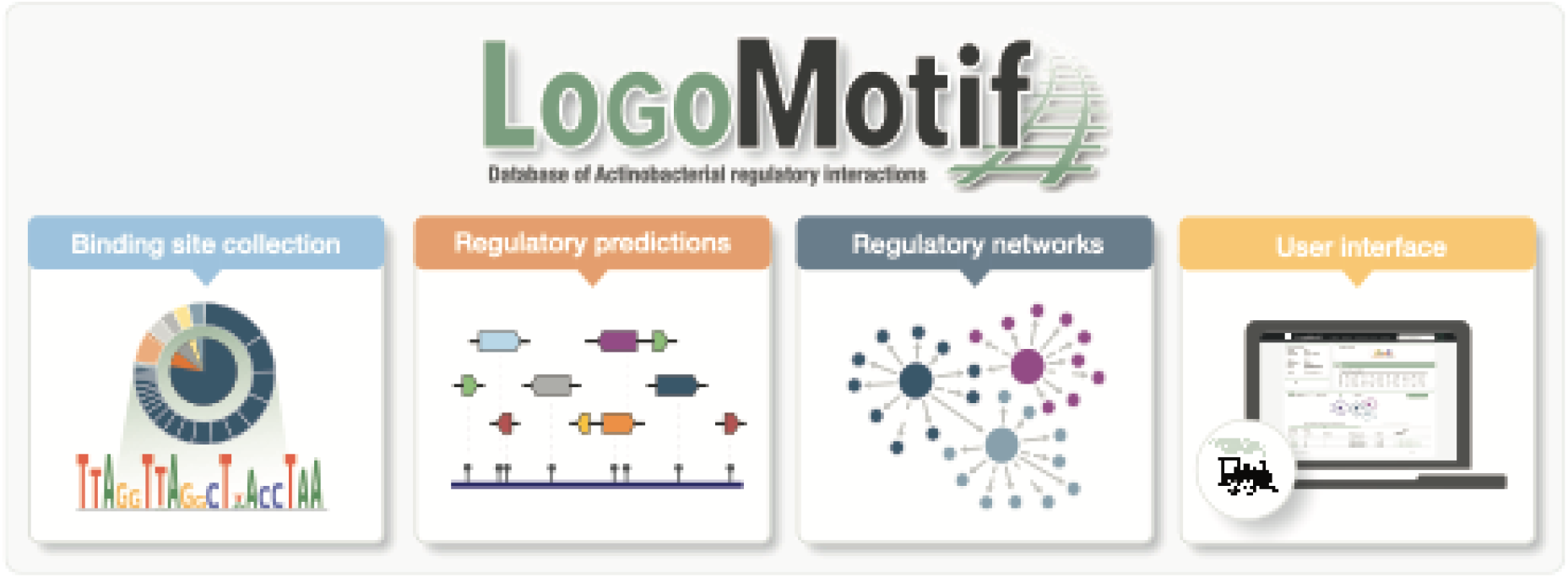

## Introduction

Actinobacteria are one of the largest bacterial phyla and known as Nature’s medicine makers [1,2]. Actinobacteria produce some two thirds of all known antibiotics and many other bioactive molecules of medical, agricultural and biotechnological importance [3,4]. Their ubiquitous presence in diverse ecosystems, both aquatic and terrestrial, necessitates their ability to rapidly perceive and respond to environmental changes [5,6]. In response to these changes, such as fluctuations in osmotic pressure, redox state, or the presence of peculiar nutrient sources, bacteria either sense or transport specific signals. These signals are either directly or indirectly linked to complex regulatory networks of multiple regulators, typically transcription factors (TFs), and their cognate TF binding sites (TFBSs), enabling bacteria to adapt to their surroundings. Together, these networks dictate the activation or repression of target genes, a process that scientists seek to understand and control, with various applications extending from strain optimization to drug discovery [7].

Insights into regulatory networks that control the biosynthesis of natural products, whose biosynthesis is encoded by biosynthetic gene clusters (BGCs), is important for drug discovery. After all, a major challenge in drug discovery is that many of the BGCs are not or poorly expressed under routine screening conditions. This is likely explained by the fact that the environmental signals that activate their expression in the habitat are missing in the laboratory [8]. In biotechnology, the challenge of low protein yields is often addressed through heterologous expression, optimizing strains and culture conditions while bypassing native regulatory systems. An example is the food industry, where polysaccharide hydrolases are typically heterologously produced in optimized production chassis for enhanced fermentation efficiency [9,10]. However, for natural products this is far less straightforward, among others for reasons of precursor supply and toxicity to the host. Therefore, expressing and optimizing pathways in the native hosts is preferred. This approach requires a comprehensive understanding of the native regulatory networks and the molecules that influence them, a crucial step for their effective characterization and application in drug discovery and various other fields.

To reliably predict how BGCs are controlled, better understanding of the binding sites and hierarchy of the TFs that control specific and global gene expression is required. Within Actinobacteria, up to 1000 TFs cooperate and antagonize each other in a multi-layered and highly complex system. The well-studied model organism *Streptomyces coelicolor* exemplifies this complexity with some 900 different regulatory proteins, of which only a small fraction has been characterized in detail [11,12]. Ironically, in the well-studied *E. coli* over 70% of the regulatory networks is known, and this was recently referred to as “ignorance” [13,14]. In *Streptomyces* only about 6% of the TF binding sites is known [15], which underlines the urgent need for more binding site data. These need to be uncovered via high-throughput methods like DNA Affinity Purification Sequencing (DAP-seq) or chromatin immunoprecipitation sequencing (ChIP-seq), and *in silico-based* methodologies [16–19]. Moreover, researchers often work with custom strains and have limited experimental data on TFBSs specific to their host of interest. To address this, computational approaches have been developed, focusing on the identification and prediction of TFBSs using models derived from experimentally validated TFBSs. Examples include PREDetector [20] and various tools of the MEME suite [21], which have proven effective in predicting TFBSs. However, actinobacterial networks have not been computed, curated and visualized in a comprehensive manner, neither in model organisms like *Streptomyces coelicolor* nor beyond.

Here, we present LogoMotif, freely accessible via https://logomotif.bioinformatics.nl/, a database of actinobacterial regulatory interactions and TFBSs. LogoMotif offers a comprehensive collection of both validated and curated genome-wide predictions of TFBSs presented in interactive regulatory networks. Additionally, LogoMotif’s collection of TFBSs serves as the foundation for the new TFBS recognition feature of the BGC prediction tool antiSMASH v7 [22]. This integration directly provides users with clues on regulatory processes in their BGCs of interest. With its continuously and actively updated collection of high-confidence actinobacterial regulatory interactions, the LogoMotif database will enable researchers to elucidate gene expression and make novel discoveries in the field of actinobacterial biology and beyond.

## Results

### A comprehensive dataset of characterized and predicted actinobacterial regulatory interactions

To compile an updated collection of TFBSs in Actinobacteria, particularly those of *Streptomyces* species, a targeted literature search was performed (Figure 1). For this search, we made use of keywords related to regulation and various experimental methods, including ChIP-seq, DAP-seq, electrophoretic mobility shift assay (EMSA), and DNase footprinting techniques. The sequences that were identified through this targeted search were manually extracted and subjected to a curation process. During this stage, we used a cut-off of at least four verified binding sites to ensure the useability of these sites for predictive modeling. This resulted in a collection of 392 experimentally characterized binding sites across 23 regulators, which in total provide approximately 15,600 predicted regulatory interactions to be explored when using default thresholds. For detailed information on the threshold setting criteria, we refer to the Methods. These interactions are visualized in interactive networks for four Streptomyces model organisms: *Streptomyces coelicolor* [23], *Streptomyces griseus* [24], *Streptomyces scabiei* [25] and *Streptomyces venezuelae* [26].

**Figure 1.**
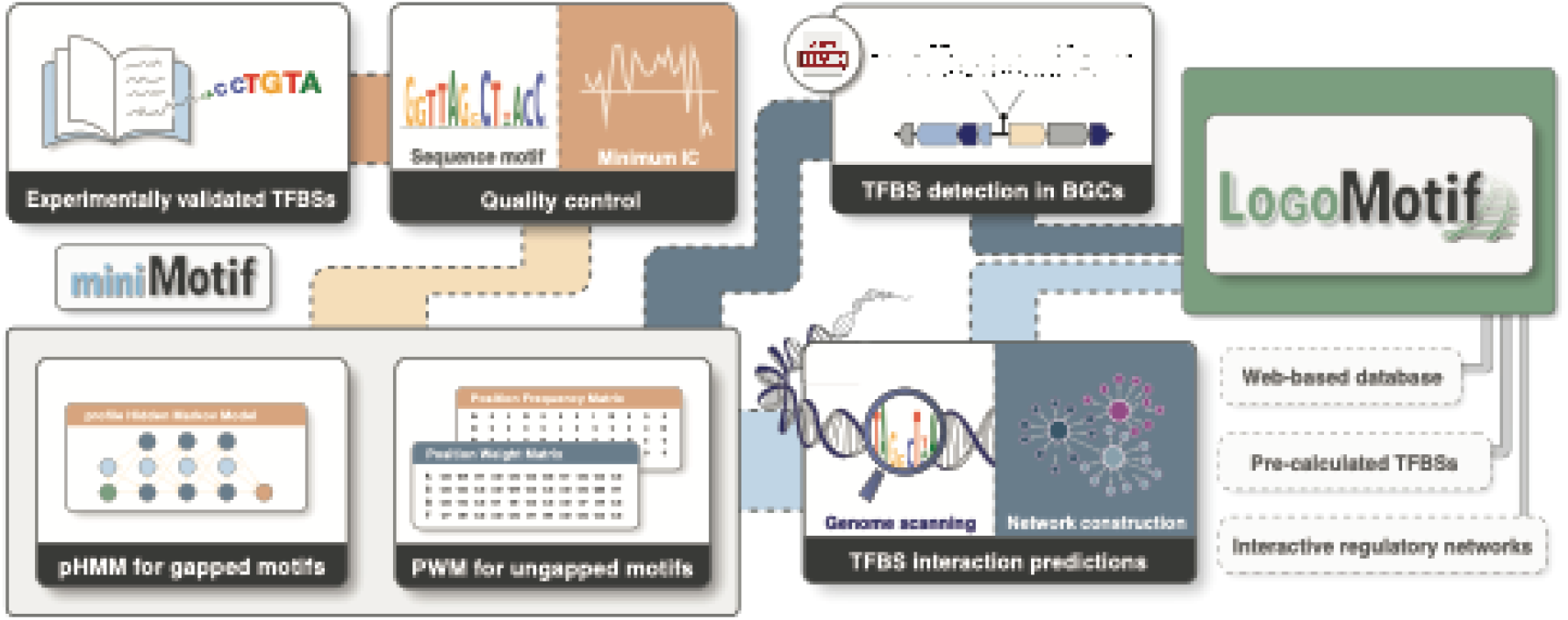
Schematic overview of the LogoMotif workflow. The process starts with a quality control step, where the information content (IC) of TFBSs gathered from literature is assessed. Depending on whether the sequences have a spacer region, either position weight matrices (PWMs) or profile hidden Markov models (pHMMs) are constructed. These models are then used to scan the genomes of various organisms, and the results are visualized as regulatory networks on the LogoMotif interface. For customized analysis, researchers can employ the MiniMotif tool (available as command-line tool at https://github.com/HAugustijn/MiniMotif/) for TFBS detection of custom strains or TFs. Additionally, the PWM detection method is integrated into the BGC prediction software antiSMASH v7, to add TFBS Finder utility. Users can access detailed regulatory information via a cross-link on the antiSMASH results page to the LogoMotif interface.

### Implementation and features of the LogoMotif database

The web interface of LogoMotif aims to offer seamless access, downloading and exploration of TFBSs for insights in regulatory interactions. At its core, the platform is powered by a SQL database, organized to store data for each regulator, including general information, sequence motifs, literature derived and predicted TFBSs, as well as regulatory networks. Upon visiting the LogoMotif homepage, a quick search feature allows for immediate querying of specific regulators. Alternatively, users can browse the catalog of regulators or follow a redirection from the TFBS Finder results in antiSMASH v7. Upon regulator selection, users are redirected to the dedicated results page, each providing in-depth insights into the chosen regulator’s data (Figure 2). The regulator page features general data of the regulatory protein itself, including cross-links to sequence, structure, and functional details in the UniProt [27], KEGG [28], and PDB [29] databases (Figure 2A & 2B). Additionally, the page displays curated binding sites as a sequence logo, along with prediction matrices and tabular data (Figure 2C, D & F). A key feature is the network visualization of both curated and predicted interactions of the regulator and its regulon (Figure 2E). This network offers users a snapshot into the regulatory cascade associated with their genes of interest, which provides insights into how genes or gene clusters may be controlled. The network contains a score threshold slider that enables users to tailor the display according to their interest. The score represents the prediction value, normalized to a maximum value of 1. This normalization is necessary to accommodate the scoring variations among different prediction models and different motif lengths. A higher score indicates a closer alignment with our model’s predictions. Obtaining a suitable threshold is important for differentiating between true and false positives, a common challenge in the detection of TFBSs [30]. Thus, users can adjust the score slider to view only the most strongly predicted sites by setting a higher threshold or choose a lower setting to explore a broader range of potential interactions. To accommodate the in-or exclusion of predictions in the network, we offer a ‘predictions’ option, which can be deselected to exclude predicted interactions from the display.

**Figure 2.**
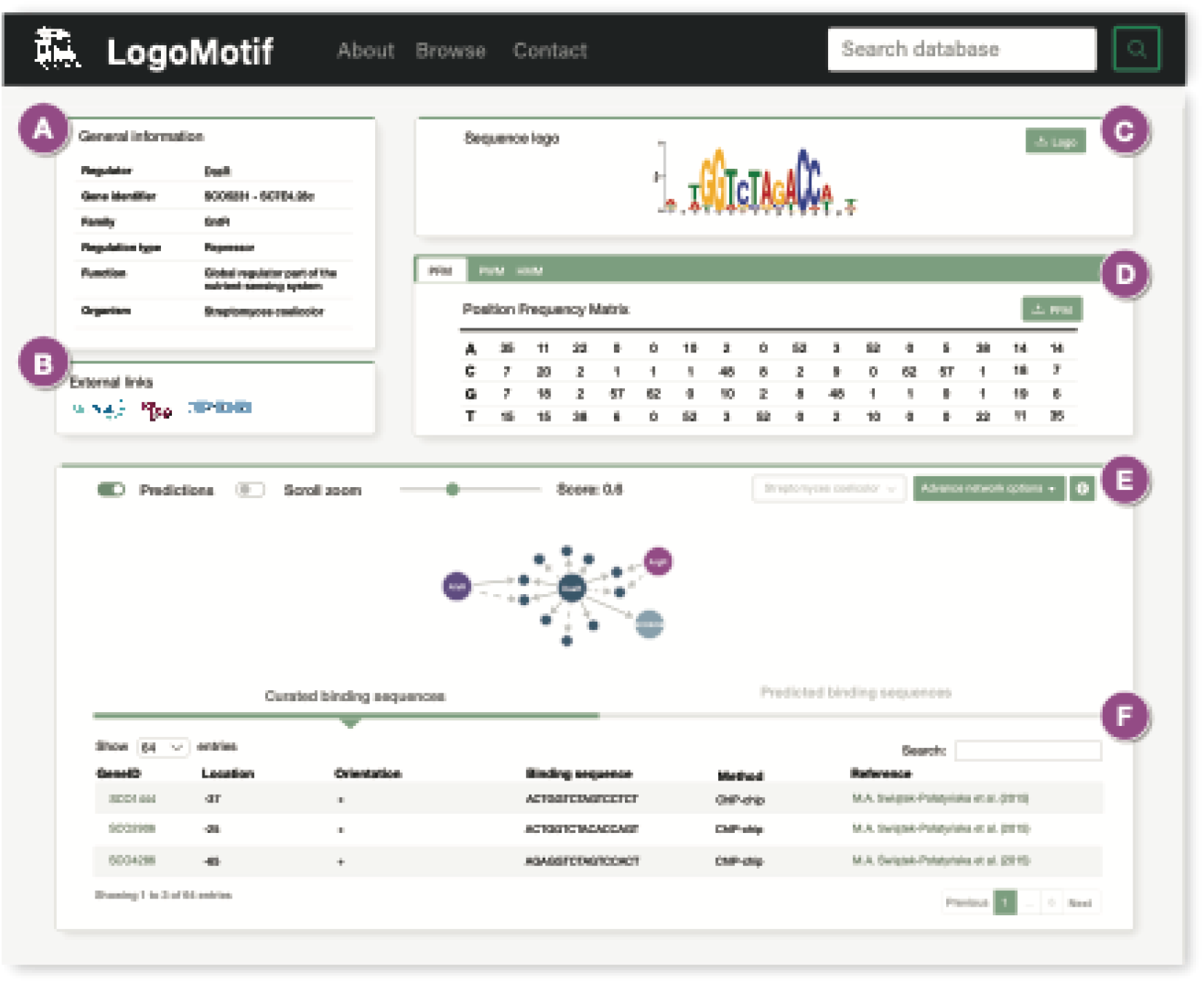
Overview of the LogoMotif user interface. **a)** General information: displays the gene name, regulatory family, and annotated functions of the regulator. **b)** Database links: provides cross-links to UniProt, KEGG, or PDB for further details on the regulatory gene, if available. **c)** Sequence logo: displays the sequence logo derived from curated binding sites, with an option to download in PNG format. **d)** Prediction algorithms: showcases the position frequency matrix (PFM), PWM, and/or HMM, specific to the regulator’s characteristics (e.g., presence or absence of spacer region). **e)** Regulatory network: shows a network of known or predicted regulatory interactions, with adjustable score thresholds to modify the network’s stringency. **f)** Binding sequences: presents both curated and predicted TFBSs.

### Models and tools for custom TFBS prediction

In addition to its collection of validated TFBSs, LogoMotif provides prediction models designed to offer deeper insights into the regulons of well-studied model strains and to facilitate regulatory research on custom strains. Users can download these prediction models directly from the LogoMotif web interface and integrate them into their own preferred analysis pipelines. Alternatively, they could use the provided TFBS prediction tool MiniMotif for fast genome wide TFBS detection on user-provided genomes or TFs. MiniMotif makes use of pre-computed position weight matrices (PWMs) and profile hidden Markov models (pHMMs) (Figure 1). With the use of PWMs, entire regulons can be predicted based on a minimal number of experimentally validated binding sites. The dual approach using pHMMs accounts for variable length spacer regions in the binding site profiles, an occurrence often found with sigma factors [31]. For each alternative spacer length, a separate pHMM is generated in which the non-conserved spacer regions are masked to improve prediction accuracy. However, in smaller datasets, PWMs are preferable to pHMMs, which are more susceptible to overfitting [32]. Both methods are applied to LogoMotif’s selection of TFs to provide easy access to pre-calculated predictions of the aforementioned *Streptomyces* model organisms. These predictions are presented as interactive visualizations on the LogoMotif interface, offering users a dynamic way to explore regulatory interactions.

### Integration and cross-links with genome mining tools

To provide insights into how silent or cryptic BGCs may be activated in the laboratory, it is of critical importance to understand the regulatory networks that control them. The recent introduction of the TFBS Finder feature in antiSMASH v7 now adds an additional layer of regulatory information (Figure 1). LogoMotif’s collection of TFBSs serves as the engine for this new feature and is based on the PWM detection algorithm of MiniMotif. In addition to this integration feature, antiSMASH users can also directly navigate to the LogoMotif webpage from the antiSMASH interface, providing them with further insights into regulatory networks and for their exploration beyond the scope of BGCs.

### Example use cases

To illustrate the different ways in which LogoMotif can be used, we provide two use case scenarios, detailing the steps that users would perform to obtain their required results.

In the first scenario (Figure 3a), an experimental scientist may be interested in investigating the regulon associated with a specific regulator to identify the range of genes directly or indirectly controlled by it, as well as their functions. Given the complexity of regulatory systems, where multiple regulators often interact and one can compensate for the loss of another, this exploration can significantly influence experimental designs. Therefore, the scientist might focus particularly on how their chosen regulator affects others. To delve into these interactions, the user can search for the regulator by name on the LogoMotif homepage. Upon finding the regulator, the user is directed to a dedicated regulator page, which includes either validated or predicted binding sites. This information is found in the lower section, under either ‘curated binding sites’ or ‘predicted binding sites’. Regulatory relationships are illustrated as a directed network, with each arrow (edge) indicating a regulatory link starting from the regulator (node) and pointing towards regulated gene nodes. To aid in the identification of interconnected regulators, regulators are visualized as larger circular nodes. Using this information, a user could gain knowledge on possible downstream effects that might occur when their regulator of interest is influenced by variables such as altered culture conditions.

In the second scenario (Figure 3b), a user employing antiSMASH for gene cluster predictions can use the TFBS Finder module to obtain insights into potential regulatory systems. The TFBS Finder identifies possible binding sites and provides preliminary information about the regulator. For a deeper understanding, the user can follow a cross-link to LogoMotif, where they can access detailed information, link to relevant literature, download motifs for visualization, and explore connected regulators through an interactive network. This enhanced overview of the regulatory system can potentially offer valuable insights into the transcription factors responsible for regulating the gene cluster. Furthermore, it can aid in the planning of additional experimental research that aims to activate the gene cluster or to uncover the underlying regulatory connections that in turn could feed the database with novel, curated TFBSs.

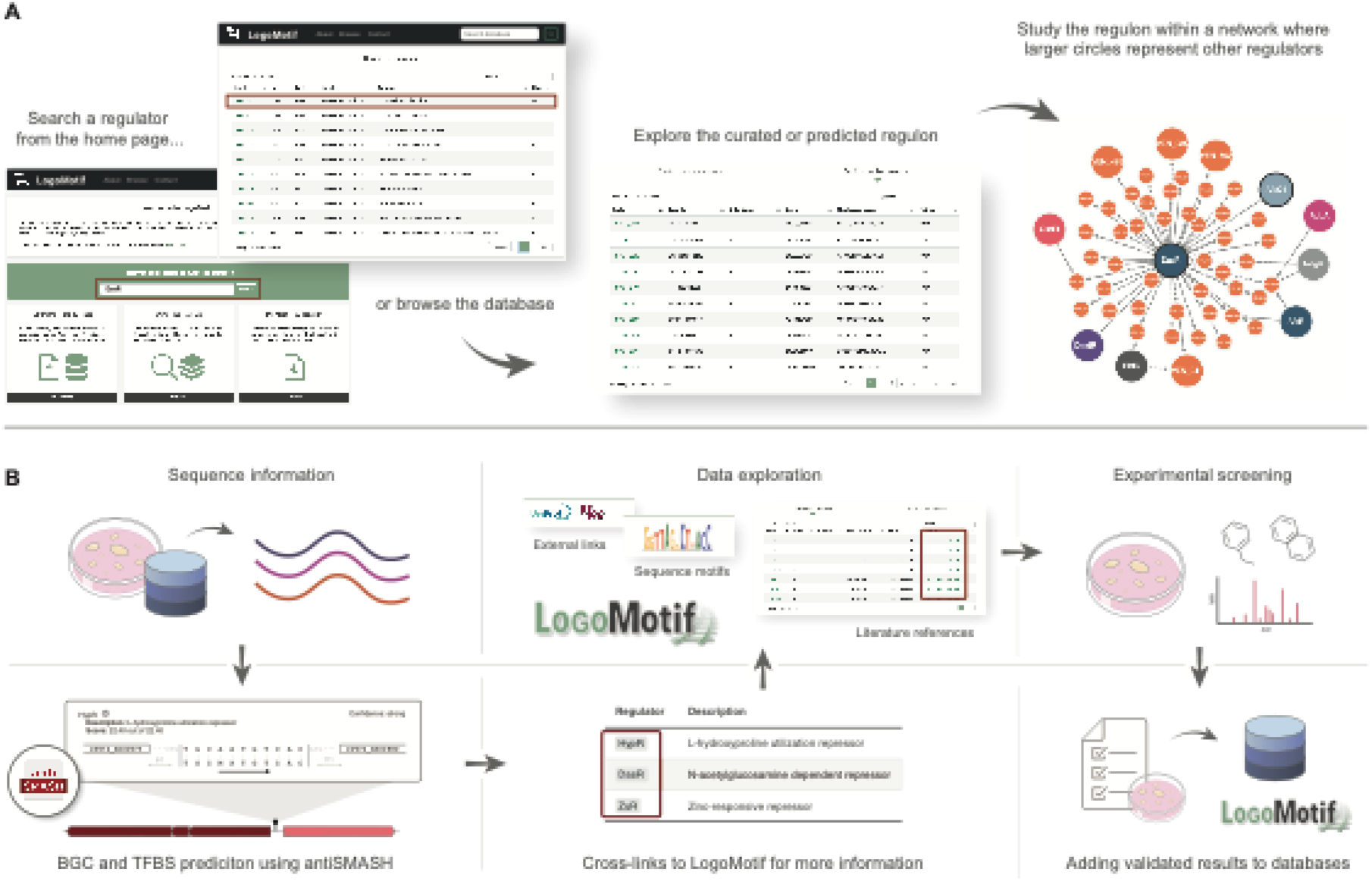

**Example use workflows. a)** Users can search for regulators via LogoMotif’s home page or through the browse page. On a regulator’s page, both curated and predicted regulons are available at the bottom, with an interactive network visualization aiding in the easy identification of regulators, highlighted by larger circles. **b)** Sequence data from experiments or databases can be inputted into BGC prediction tool antiSMASH v7, which offers regulatory predictions via the TFBS Finder and provides cross-links to LogoMotif. The LogoMotif page further offers links to literature, related databases, and provides sequence logos for TF visualization purposes. This information aids in hypothesis generation and gives leads for experimental validation, with the option to add new findings to the database, improving its knowledge base.

## Discussion

LogoMotif is a new database that focuses on providing insights into the regulatory interactions in members of the Actinobacteria, one of the largest bacterial phyla. Thus, LogoMotif complements databases such as Prodoric [33] or RegPrecise [34], which focus primarily on Firmicutes and Proteobacteria, respectively. The LogoMotif database is not merely a collection of TFBSs retrieved from scientific literature, but also a platform for genome-wide predictions across four *Streptomyces* model organisms. LogoMotif distinguishes itself from other databases by using a combination of PWMs and HMMs for enhanced prediction of binding sites, particularly those with variable length spacer regions. This approach is provided to the user via MiniMotif, a command-line tool that offers researchers exploring regulatory interactions of custom strains or TFs. Additionally, LogoMotif’s collection forms the base of the TFBS Finder module integrated within antiSMASH v7, providing TFBS predictions for research specifically interested in BGC regulation. The curated and predicted TFBSs are presented in interactive regulatory networks, enabling researchers to delve deep into the dynamics of Actinobacterial gene regulation.

Despite the current collection of TFs in LogoMotif being highly valuable, it represents only a fraction of the complete regulatory landscape of Actinobacteria. The experimental characterization of binding sites is a challenging and time-consuming task. Traditional methods, such as EMSAs, offer limited throughput and the reported binding interactions, frequently presented in figures or not fully made publicly available, are difficult to extract and incorporate in modern databases. However, in the postgenomic era the field is changing rapidly with the introduction of cost-effective, high-throughput experimental methods, promising an increase in the availability of large, curated datasets. Therefore, we anticipate a large increase in our knowledge base in the coming years. LogoMotif is designed to accommodate this growth and will serve as an open science hub to incorporate and harness this information for the scientific community at large.

In following releases, the LogoMotif interface will be updated with new releases, which will include a genome browser, enabling users to visualize binding sites within their genomic context, and integration of MiniMotif directly into the website interface, thus facilitating easier access and utilization. Moreover, we are currently expanding the knowledge of the known TFBSs via large-scale DAP-seq experiments, which is expected to enlarge the repository of regulatory interactions and prediction models by 5-to 10-fold. These advancements will allow users to delve deeper into the regulatory systems governing their genes of interest, offering insights into possible triggers for gene expression. With the current collection, and those we see on the horizon, we aim to provide grounds for regulatory discoveries and subsequent utilization across numerous research domains.

## Materials and Methods

### Data curation

On the literature collected TFBSs, we identified the corresponding sequence motifs using MEME v5.5.4 [21] of each individual TF. Additionally, we performed an additional manual curation step if the sum of Shannon’s entropy information content (IC) scores across all positions within the motif were less than half of the maximum IC score relative to its length, to ensure that the motif was not the result of random noise. This step involved a re-calculation and re-evaluation of the IC scores and motifs to confirm the reliability and accuracy of our motifs.

### Construction of computational prediction models

The back-end system of LogoMotif is combined into a python-based command line package named MiniMotif (accessible via https://github.com/HAugustijn/MiniMotif/). This pipeline makes use of MEME v5.5.4 [21] for motif discovery, Logomaker v0.8 [35] for the visual representation of the motifs, Bioconductor’s seqLogo v5.29.8 [36] for the construction of PFMs and PWMs, and HMMER v3.3.2 (http://hmmer.org/) for the construction of pHMM profiles. The genome assemblies of four model organisms were downloaded from NCBI in Genbank format. This includes *Streptomyces coelicolor* A3(2) (GCA_000203835.1), *Streptomyces griseus* subsp. *griseus* NBRC 13350 (GCF_000010605.1), *Streptomyces scabiei* 87.22 (GCA_000091305.1) and *Streptomyces venezuelae* ATCC 10712 (GCF_000253235.1). Based on the principles of PREDetector [20], on default, the region -350 bp to +50 bp relative to the start codons of each gene of these genomes were extracted with the use of MiniMotif. MOODS v1.9.4.1 [37] was used with a p-value of 1 × 10^−5^ cutoff to query these regions for the presence of matches to PWM matches. Additionally, the default network threshold for the PWM is determined by summing the positions with an IC score exceeding half of the max IC score, which aids in distinguishing stronger matches to the PWM. All matches detected from the p-value threshold and onwards are reported to offer a comprehensive overview of potential interactions. For the HMM profiles, input sequences were aligned using MAFFT v7.52 [38], whereafter a background frequency distribution was assigned to nucleotides belonging to non-conserved spacer regions using HMMER alimask. Next, nhmmscan was used for TFBS detection with a 0.1 bitscore threshold and a filtering step is performed to remove partially aligned hits that only cover a fraction of the pHMM. Only sequences that align with the pHMM and that exceed this threshold are reported in the final output.

### Web application implementation

The LogoMotif web application was developed using a python Flask framework (https://palletsprojects.com/p/flask/) for request handling and server-side routing. For the user interface layout, we employed Bootstrap v5.1.3 (https://getbootstrap.com/docs/5.1) and custom stylesheets to complement Bootstrap’s base styling. For data storage, we integrated a PostgreSQL database (https://www.postgresql.org/) and used SQLAlchemy (https://www.sqlalchemy.org/) to manage the interaction between our python code and database. Visualization of regulatory networks was achieved using the JavaScript library cytoscape.js (https://js.cytoscape.org/).

### Code and data availability

LogoMotif is freely available at https://logomotif.bioinformatics.nl/. Novel or newly submitted TFBSs will be made available with regular updates. The code for binding site processing is available via MiniMotif (https://github.com/HAugustijn/MiniMotif/). Both the web-interface and underlying code will be regularly maintained.

### CRediT authorship contribution statement

**Hannah E. Augustijn:** Conceptualization, Methodology, Software, Data curation, Visualization, Writing - original draft, Writing - Review & Editing. **Dimitris Karapliafis:** Methodology, Software, Data curation, Writing - Review & Editing. **Kristy Joosten:** Software, Writing - Review & Editing. **Sébastien Rigali:** Data curation, Writing - Review & Editing. **Gilles P. van Wezel:** Conceptualization, Supervision, Writing - original draft, Writing - review & editing. **Marnix H. Medema:** Conceptualization, Supervision, Writing - original draft, Writing - review & editing.

## Acknowledgements

The work was supported by the European Union via ERC Advanced Grant 101055020-COMMUNITY to G.P.v.W. and ERC Starting Grant 948770-DECIPHER to M.H.M.

